# Detection of Base Analogs Incorporated During DNA Replication by Nanopore Sequencing

**DOI:** 10.1101/549220

**Authors:** Daniela Georgieva, Qian Liu, Kai Wang, Dieter Egli

## Abstract

DNA synthesis is a fundamental requirement for cell proliferation and DNA repair, but no tools exist to identify the location, direction and speed of replication forks with base pair resolution. Mammalian cells have the ability to incorporate thymidine analogs along with the natural A, T, G and C bases during DNA synthesis, which allows for labelling of replicating or repaired DNA. The Oxford Nanopore Technologies (ONT) MinION infers nucleotide identity from changes in the ionic current as DNA strands are pulled through nanopores and can differentiate noncanonical nucleotides from natural ones. Here, we demonstrate the use of the ONT MinION to detect 11 different thymidine analogs including CldU, BrdU, IdU, as well as, EdU alone or coupled to Biotin and other bulky adducts in synthetic DNA templates. We also show detection of IdU label, incorporated during DNA replication in the genome of mouse pluripotent stem cells. We find that different modifications generate variable shifts in signals, providing a method of using analog combinations to identify the location and direction of DNA synthesis and repair at high resolution. We conclude that this novel method has the potential for single-base, genome-wide examination of DNA replication in stem cell differentiation or cell transformation.

## INTRODUCTION

DNA replication is a fundamental requirement for the development of an organism and the life-long maintenance of organ function. An estimated 2-3×10^11^ cell divisions occur in the human body each day to replenish lost cells and repair tissue damage. While DNA is replicated just once during a cell cycle, the pattern by which this occurs varies between cell types [1], and is different in malignancy [2]. The pluripotent state, for example, is maintained by a replication program very different from the one in mature cells [3]. Thus, during the reprogramming of a somatic cell to a pluripotent stem cell, the replication timing pattern is reset to resemble the one in embryonic stem cells [4]. Importantly, cancer cells also acquire replication programs which differ from the ones in their normal counterparts. For instance, allelic replication asynchrony at the p53 and 21q22 loci has been described in invasive carcinomas [5]. Delayed replication of one allele of a tumor suppressor can interfere with gene expression and create a situation similar to loss of heterozygosity which is associated with malignancy [5]. Chromosome-wide delays in DNA replication have been shown to delay chromosome condensation and contribute to the chromosomal instability of various tumor cell lines [6]. Therefore, in addition to studying differential gene expression patterns, there is a strong rationale to study differences in DNA replication patterns in development and disease.

Methods that are currently available to analyze the progression of DNA replication include a pulse of BrdU and immune-precipitation for labelled DNA [7]. Replication patterns can also be inferred from the number of reads in next generation sequencing data [8], as well as from sequencing of Okazaki fragments [9]. Next generation sequencing allows the analysis of DNA replication timing, and can identify the location of replication origins. However, it requires amplification during library preparation and averages the signal from different cells and DNA strands. To identify the progression of replication on a single DNA strand, a commonly used technique is DNA combing, which relies on sequential pulses with two modified nucleotides-IdU and CldU, fiber stretching on glass slides and staining with specific antibodies. While this technique can track replisome progression, stalling and restart in a genome-wide manner, it does not provide any sequence information. To study replication dynamics at a locus of interest, fiber analysis requires combination with fluorescent in situ hybridization [10]. This method, however, is not scalable to the level of a complete eukaryotic genome. Thus, to the best of our knowledge, there is currently no technique that can capture location, direction and speed of replication fork progression on single molecules at a genome-wide level with single-base resolution.

Nanopore technology has recently emerged as a powerful third-generation sequencing technique, primarily utilized for genome assembly due to its ability to produce ultra-long reads [11]. The system uses electrophoresis to transport DNA through a collection of nanopores and bases are identified by measuring changes in the electric current across the surface of the orifice [12]. In reality, signal shift is produced not by a single base, but is influenced by 5-6 neighboring nucleotides, making contact with the pore and termed a k-mer. Computational algorithms are then used to process the signal and determine the identity of the bases passing through. Since the nanopore platform does not require DNA amplification for library preparation, it can sequence cellular DNA directly and thereby avoid averaging signals from different DNA strands. Because nanopore technology does not rely on base-pairing to determine DNA sequence, it may be used to distinguish not only the four canonical nucleotides (A, T, G, C), but potentially other types of base. Indeed, the system is able to detect methylated bases [13], which suggests that the sequencing of nucleotide analogs incorporated during DNA replication may also be possible.

Here, we present the use of nanopore sequencing on the Oxford Nanopore Technologies (ONT) MinION to detect thymidine analogs that can be directly incorporated during DNA replication or generated in click cycloaddition reactions. Using a series of templates, we show that the MinION can distinguish between thymidine and all 11 analogs, with the greatest signal to noise ratio shown by IdU, CldU and Biotin-dU. We also show detection of IdU in substituted mammalian DNA. The methods presented in this study provide a path towards extracting comprehensive, genome-wide information on the origin, direction and progression of polymerases during DNA replication and repair from MinION sequencing reads.

## MATERIAL AND METHODS

### Training Template Assembly

For training sequence assembly, the pCALNL-GFP vector (Addgene plasmid#13770) was digested with AgeI and NotI to release the E-GFP insert. A double stranded DNA oligomer with 5 nbBbvCI sites was prepared by hybridizing two single-stranded DNA oligos and ligated into the purified empty vector as described in Luzzietti et al.[14]. The product was transformed into E-coli. Nicking and substitution reactions were performed as outlined in [14] with oligos, purchased from Baseclick, Germany. Prior to MinION library prep, all templates were linearized with SpeI (New England Biolabs).

Top Clone:

5’-CCGGGCCTCAGCTTGCCACGACCTCAGCGTCAATTGTCCTCAGCTCAGATGACCTCAGC AGATTGTAGCCTCAGC-3’

Bottom Clone:

5’-GGCCGCTGAGGCTACAATCTGCTGAGGTCATCTGAGCTGAGGACAATTGACGCTGAGGT CGTGGCAAGCTGAGGC-3’

Replace Oligo Single:

5’Phos-TGAGGCTACAATCTGCTGAGGTCATCTGAGC**X**GAGGACAATTGACGCTGAGGTCGT GGCAAGC-3’ X = EdU, CldU, BrdU, IdU

Replace Oligo Multi:

5’Phos-**X**GAGGC**X**ACAA**X**C**X**GAGG**X**CA**X**C**X**GAGC**X**GAGGACAA**XX**GACGC**X**GAGG**X**CG**X**G GCAAGC-3’ X = EdU, CldU, IdU

### Click Cycloaddition Reactions

Click cyclo-addition was performed on single stranded EdU replace oligos prior to the substitution reaction. Briefly, 1ug phosphorylated oligo was incubated with reagents from the CuAAC Biomolecule Reaction Buffer kit (Jena Bioscience CLK-072) and 250uM azide for 1h at 37C. Separate reactions were carried out with the following azides: sodium azide (Sigma S2002), Nicotinoyl azide (Sigma CDS006775), 6-azido-6-deoxy-D-glucose (Sigma 712760) Biotin azide (Thermo Fisher B10184), AF488 azide (Thermo Fisher A10266), AF647 azide (Thermo Fisher A10277), and a ssDNA oligo purchased from Integrated DNA Technologies: 5’-GGATAGCCTC/3AzideN/-3’. All click products were purified by precipitation.

### Dot Blots

For dot blots, 100-200ng DNA was denatured with 20mM NaOH for 5min at 99C, cooled on ice and neutralized with 0.6M ammonium acetate. Spotting was performed manually on a 0.45uM nylon membrane (Thermo Fisher 77016). Once fully dried, the membrane was baked in a microwave for 2.5min. Prior to imaging, Biotin-containing blots were stained with A647-Streptavidin (Thermo Fisher S32357). For this procedure, the baked membrane was incubated with 3% BSA in TBST for 15min, followed by A647 Streptavidin 1:500 (Thermo Fisher S32357) for 45min at room temperature and 3 washes with TBST. To detect CldU or IdU-containing DNA, 500ng-1ug of sample was spotted on the membrane and stained with rat BrdU/CldU 1:500 (Bio Rad OBT0030S) or mouse BrdU/IdU 1:500 (BD 347580) in 3% BSA in TBST for 1h at room temperature. Staining with mouse ssDNA antibody 1:500 (Millipore MAB3034) was used as a loading control. To visualize the signal, the membrane was incubated with Alexa Fluor 647 and Alexa Fluro 488 secondary antibodies at 1:500. Following washes, the membranes were imaged with a Biorad ChemiDoc MP Imaging System.

### MinION Runs on Synthetic Templates

All samples were sequenced on MinION R9 flow cells. Sequencing libraries were prepared with Ligation Sequencing Kit 1D (ONT SQK-LSK108), following manufacturer’s instructions. Each run was performed with 250-500 ng purified DNA. Typically, a single flow cell was used for multiple samples with washes in between, carried out with the Flow Cell Wash Kit (ONT EXP-WSH002). Fast5 data files from each run were analyzed with NanoMod as outlined in Liu et. al [17].

### MinION Runs with Genomic DNA

For genomic DNA runs, mouse embryonic stem cells from a pure C57BL/6J strain (Jackson LAB Strain # 000664) were incubated with 25uM IdU for 24h. Cells were then harvested and genomic DNA was extracted with a High Pure PCR Template Preparation Kit (Roche 11796828001). MinION runs were performed on R9 with 400-500ng purified DNA per sample following library preparation with Ligation Sequencing Kit 1D (ONT SQK-LSK108). For each run, unlabelled DNA was loaded first and sequenced for 5-6 hours. This was followed by a wash step with the ONT wash kit (ONT EXP-WSH002), priming and loading of IdU-substituted gDNA which was run until the flow cell expired, typically from 8 to 20 hours without voltage adjustments. Analysis was performed with data from two combined runs. The combined control data set included 719,295 long reads covering ∼2.4G bases while the IdU sample was represented by 140,667 long reads across 417M bases.

## RESULTS

### Detection of single-base DNA modifications with nanopore sequencing

To determine the ability of the ONT MinION sequencer to detect modified bases, we designed synthetic templates containing a single modification in a defined location. Template assembly involved a vector nicking step with nb.BbvCI which generated a 63bp region of ssDNA, followed by the ligation of an oligo with one modified thymidine base (Fig. 1A). This approach to replace segments within a plasmid with a labelled oligo has previously been used for FRET studies and reported to result in >75% substitution [14]. We used oligos without modification, or containing 5 different modifications, EdU, CldU, BrdU, IdU, or Biotin for the replace reaction (Fig. 1B). Also, we modified EdU with a series of compounds of increasing molecular weight: azide, nicotine, glucose, AF488, AF647 and a 10bp single stranded DNA oligo through click cycloaddition reaction [15]. This method allows the attachment of any azide group containing moiety to alkyne-substituted DNA and has been reported to be nearly 100% efficient [16]. Dot blots for AF647, AF488, BrdU and Biotin modified plasmids showed efficient labelling (Fig. 1C). Following the replace reaction, the vector was linearized with SpeI to generate a 6184-bp double-stranded training template with one modified thymidine on the minus strand at position 3073, which was sequenced on the MinION (Fig. 1D).

**Figure 1.**
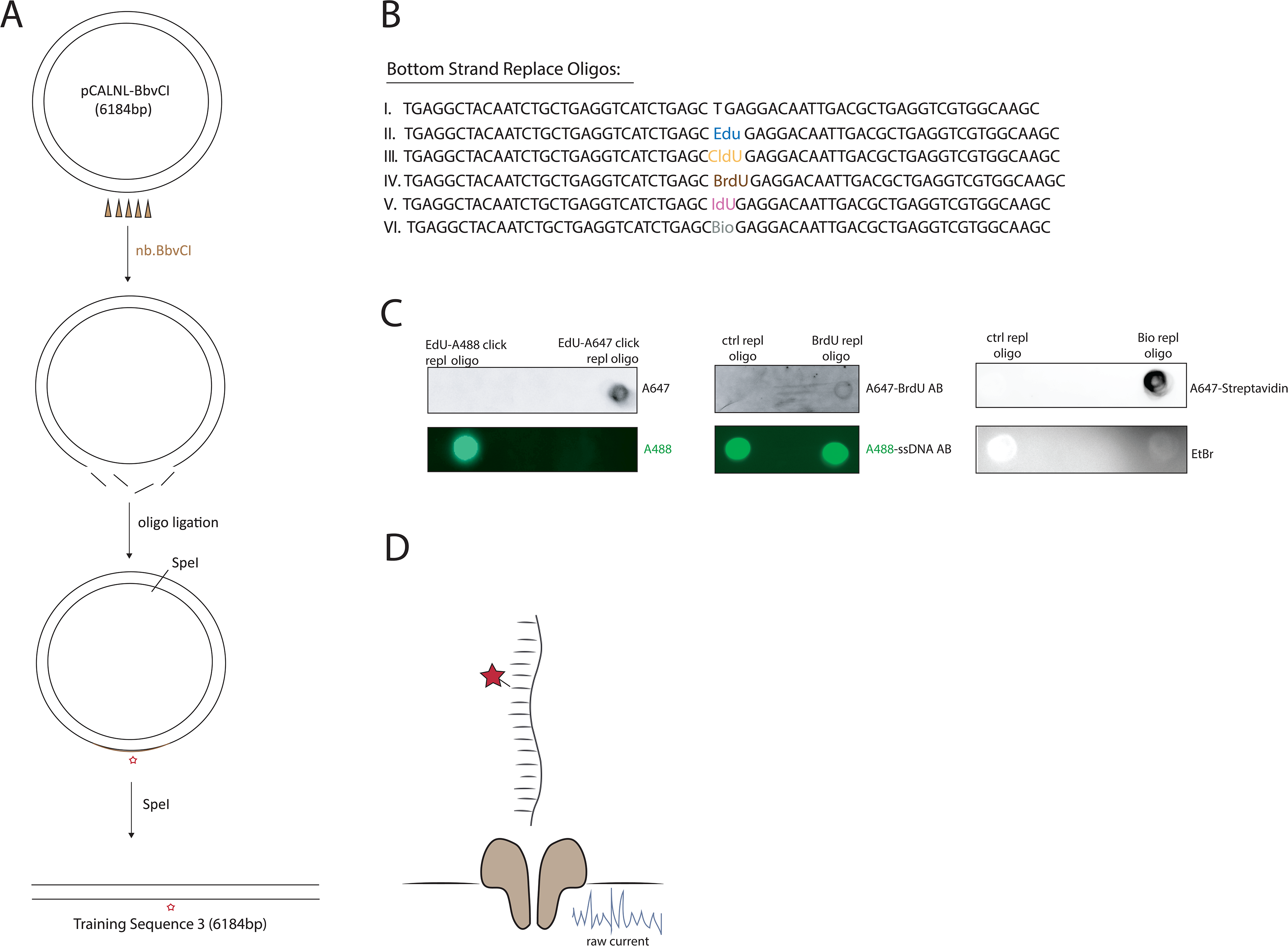
Design and assembly of training templates with single modifications. A). A schematic of the two-step assembly of single modification training templates. The first step is nicking with nb.BbvCI and the second-ligation of a ssDNA oligo which carries a modification. The vector is then linearized with SpeI to generate a 6184bp sequence with one thymidine analog at position 3073. B). Sequence of the single modification oligos used for ligation. C). Dot blots with ligation products for AF647, AF488, BrdU, IdU and Biotin modifications, demonstrating efficient replacement with the modified oligo. D). A schematic illustrating the sequencing of a training template with a single-modification through a nanopore.

To identify modified nucleotides, we developed NanoMod, an analysis tool which calls DNA modifications directly from electrical signals [17]. NanoMod uses Albacore for base calling and performs indel error correction by aligning signals to a reference sequence. The input to NanoMod is a dataset with two groups of samples-a modified template and a sequence-matched control, while the output is the ranked list of regions with modifications. Signals from control and substituted samples are then compared using two statistical assays-Komogorov-Smirnov test and Stouffer’s method for calculating combined p values. We used NanoMod to score individual nucleotides in 5-bp sliding windows to detect the presence of thymidine analogs. All 11 modifications (Fig. 2A) in the dataset caused signal shifts at base 3073 as well as at neighboring positions (Fig. 2B-M). The analogs EdU, CldU, BrdU and IdU caused signal shifts spreading as far as 3-4 bases upstream and 2 bases downstream of the modified position (Fig. 2B, C, D, F), consistent with the minion detecting signals from a 5-6 bp k-mer. Therefore, the signal of the modified T is present in neighboring bases. The magnitude of the signal change was proportional to the size of the moiety with one of the largest unclicked analogs IdU (MW=354.1g/mol) generating the most significant change in pore current, log(Combined p-value) = −175 (Fig. 2F). The smallest modification, EdU (MW=252.23 g/mol) was detectable as well at log(Combined p-value) = −125 (Fig. 2A). A highly significant change in signal at 3073 was also recorded for the bulkier analog-Biotin, with a log(Combined p-value)=-150 (Fig.2I). Purification of the plasmid after the replace reaction with streptavidin beads, followed by sequencing further increased the significance of the change, with a log(Combined p-value) = −250 (Fig. 2J). Therefore, modification at 75% [14], was sufficient to call the modified base, while 100% modification further magnified signal change. Biotin influenced electrical signals further away from the modification than EdU, CldU, BrdU or IdU. An increased number of surrounding bases affected-as many as 3-4 nucleotides upstream and about 10 downstream of the modified position (Fig. 2I, J) indicates that the effect of bulky modifications can spread beyond a single k-mer.

**Figure 2.**
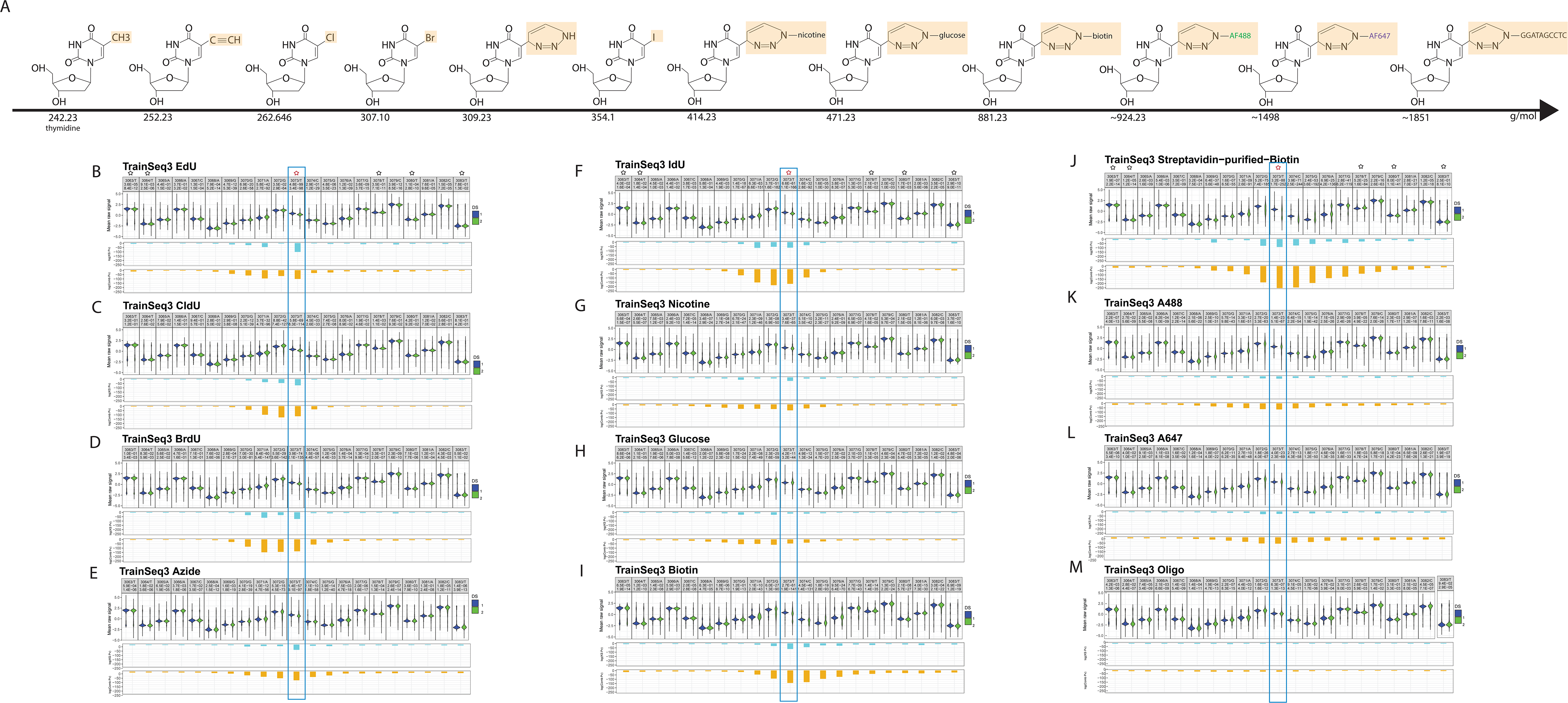
Detecting a series of single-base modifications with increasing molecular weight through MinION sequencing. A). Chemical structure and molecular weight of the tested analogs. B). A violin plot of the known modification site and the surrounding bases for B). EdU, C). CldU, D). BrdU, E). Azide, F). IdU, G). Nicotine, H). Glucose, I). Biotin, J). Streptavidin purified Biotin, K). AF488, L). AF647 and M). a 10bp ssDNA oligo, attached to position 3073 by click cycloaddition. In each panel, the first line denotes position, followed by the base, the second line shows mean raw signal from the control and modified sample, the third line contains the p-value (pv for short) for a Kolmogorov-Smirnov test, and the fourth line displays the combined p-value, calculated using Stouffer’s method. ‘DS 1’ in blue represents the non-modified oligo samples, while ‘DS 2’in green stands for the sample with a modification at position 3073. ‘-strand’ denotes reverse strand. Black stars represent unmodified T’s and the red star marks the modified position.

We also compared modifications attached to the training template by click cycloaddition - an azide, nicotine, glucose, AF488 and AF647. In regions outside the modified base, we did not observe significant changes in signals, indicating that the click reaction did not affect the ability to sequence DNA on the MinION (e.g. compare position T3063 and T3083 on the control and experimental templates in Fig. 2G). At the modified base, nicotine showed a significant change with a log(Combined p-value) = −74 (Fig. 2G). The mean raw signal of bulky bases caused a wider spread in signal, which generally affected 5-6 bases preceding the modification and about 2-4 bases downstream (Fig. 2E, G, H). Clicked moieties AF488 and AF647 caused a more extensive downstream current shift, spreading as far as 8-10 bases beyond the modified position (Fig. 2K, L). The mean raw signal at the modified base was visibly different when compared to control, though the change did not translate into lower p-values in the NanoMod statistical analysis. The largest clicked modification - a ssDNA oligo, expected to generate a branched structure, caused the smallest change in signal which spread over 10 bases upstream and 2-3 positions downstream of 3073 (Fig. 2M). In this particular case, reduced progression through the pore, or signal integration from bases on the training template and the clicked oligo might have contributed to the low shift in current recorded for the modified position.

These observations point to a relationship between adduct size and the magnitude of signal shift at the modified position as well as the neighboring bases. In general, bulkier moieties caused more pronounced changes in the raw signal at the modified position, and affected more bases in its vicinity. The differences in signal spread between adducts of different molecular weight show that pairs of analogs, such as IdU and Biotin, can be distinguished from each other based on the width of the window affected by nanopore current shifts (Fig. 2F and Fig. 2J).

### Detection of multiple DNA modifications with nanopore sequencing

To mimic the scenario of pulsing replicating cells with nucleotide analogs, which are incorporated at multiple positions into newly synthetized DNA, we repaired the nb.BbvCI nicks on the minus strand of the training sequence by ligating an oligo with 14 thymidine modifications between positions 3042 and 3104 (Fig. 3A). Based on the results with a single defined position, we selected 4 analogs - EdU, CldU, IdU, and Biotin added to EdU by click cyclo-addition (Fig. 3B). Efficient replacement was confirmed by dot blots for CldU, IdU and clicked Biotin (Fig. 3C). Templates with replacements were sequenced and analysis was carried out by dividing the reference sequence into ∼400 regions of 60bp each with a 30-bp overlap between adjacent windows. For all analogs tested, NanoMod correctly identified the modified region between 3042 and 3104 as shown by the p-values for Kolmogorov-Smirnov test and Stouffer’s method (Fig. 4A-D). The most significant shift of signal was centered around a modified base, gradually abated with increasing distance from the analog and raised again in the vicinity of the next modification on the template. Within each template, some modified thymidines generated stronger signals than others (Fig. 4A-C). As in templates with single modifications, the magnitude of signal shift in sequences with multiple analogs was determined by the size of the adducts with IdU generating the greatest signal to noise ratio (Fig. 4C). Biotin, the largest moiety, caused signal changes spreading over the whole region between 3042 and 3104. This spread of signal is likely related to the chemical structure of biotin, and the ability of bulky adducts to influence the current reading beyond a single k-mer. This dataset on multiple modifications demonstrates that MinION sequencing and analysis with NanoMod can be successfully used to identify short genomic regions with modified bases in DNA.

**Figure 3.**
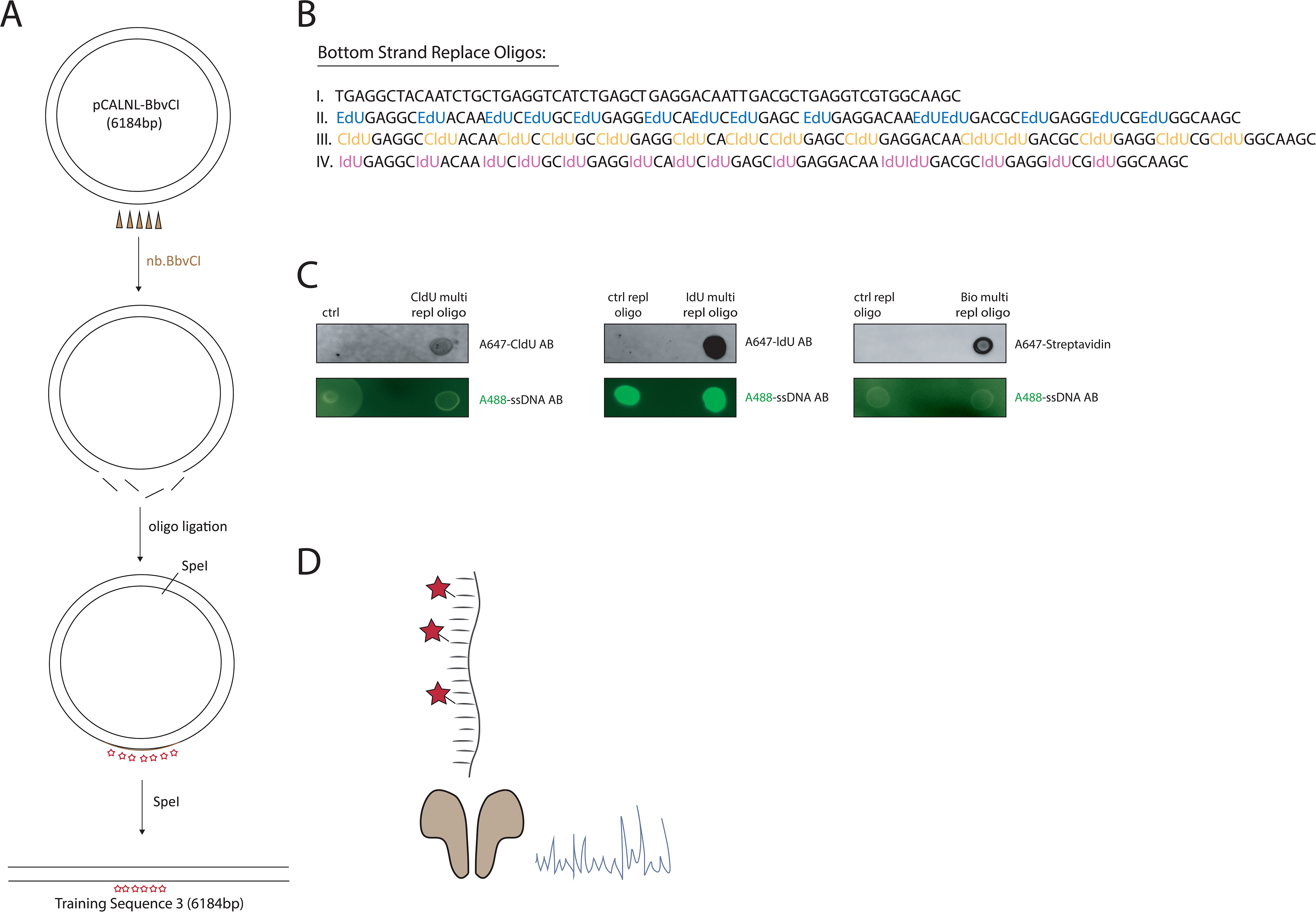
Design and assembly of training templates with multiple modifications A). A schematic of the strategy used to assemble training templates with multiple modifications. A ssDNA oligo carrying 14 analogs is used in the ligation step. B). Sequence of the multiple modification oligos used for ligation. C). Dot blots with ligation products for CldU, IdU and Biotin. CldU and IdU were visualized by antibody staining on the blot. Biotin was seen after incubation with Alexa 647-streptavidin. D). An illustration of sequencing a DNA template with multiple analogs through a nanopore.

**Figure 4.**
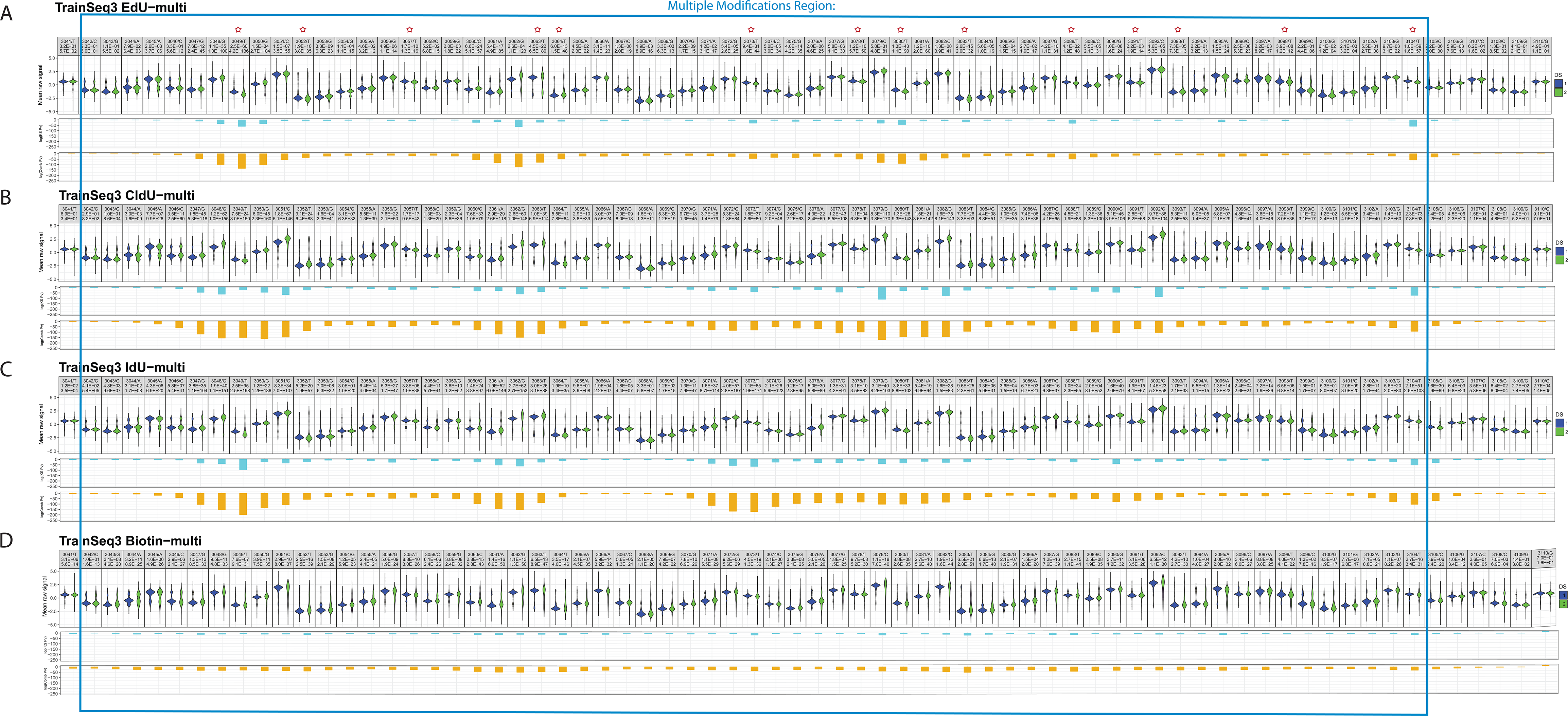
Detection of multiple modifications with MinION sequencing. A) A violin plot of the modified region 3042-3104 and surrounding bases for EdU, B). CldU, C). IdU, and D). Biotin clicked to an EdU multi modified template. ‘DS 1’ in blue represents the non-modified oligo sample, while ‘DS 2’ in green stands for the multi modified template. The Kolmogorov-Smirnov test p-value is shown on line 3, while the combined p-value, calculated by Stouffer’s method is displayed in line 4. Red stars indicate the position of modified T’s.

### A comparative analysis of nucleotide analogs for gDNA modifications

To determine if pore sequencing and the analysis method NanoMod developed here can be applied to sequencing base analogs incorporated by replicating cells in genomic DNA, we performed computational simulations. We first determined the level of analog substitution *in vivo* by exposing primary human fibroblasts to EdU for 24h, or a complete cell cycle. Purified, EdU-containing genomic DNA was then labelled with AF647 in a click cycloaddition reaction and the level of EdU-AF647 substitution was determined by measuring sample fluorescence against a standard curve of PCR products with known percentage substitutions of EdU, clicked with AF647. This analysis revealed that 20% of thymidine bases in genomic DNA were substituted with EdU (Supplementary Fig. 1A, B). This is the expected percent modification for replication tracts pulsed with thymidine analogs[18]. Based on this result, we performed computational simulations with 20% modified and 80% unmodified reads to determine what coverage would be required to detect single (Fig. 5A) and multiple (Fig. 5B) modifications in genomic DNA. Since the fraction of modified reads without purification is approximately 75% [14], the proportion of modified reads in the simulation is approximately 15%. IdU and CldU were easiest to detect in the group of analogs that can be directly incorporated during DNA synthesis, requiring 450 reads of a modified site to rank strands with a single modification in the 0.05 percentile or less (Fig. 5A). The calling of BrdU, CldU and the calling of single EdU-modified strands also obtained the highest 0.05 percentile rank as low as 450 or 950. Similarly, Biotin modifications were ranked in the 0.05 percentile range with only 450 reads. The clicked analogs-an azide group, nicotine and glucose required 950 reads, almost twice the number noted for IdU and CldU, to achieve the 0.05 percentile rank. The clicked fluorophores AF488 and AF647 required more extensive coverage-1950 and 2450 reads, respectively for more than 75% of the strands to be ranked in the 0.05 percentile or between percentiles 0.05 and 0.12. This result is consistent with the wide-spread signals of lower significance described for AF488, AF647 and the ssDNA oligo in Fig. 2. This result corroborates the observation that thymidine analogs of higher molecular weight IdU, and Biotin are readily detected on the MinION platform.

**Figure 5.**
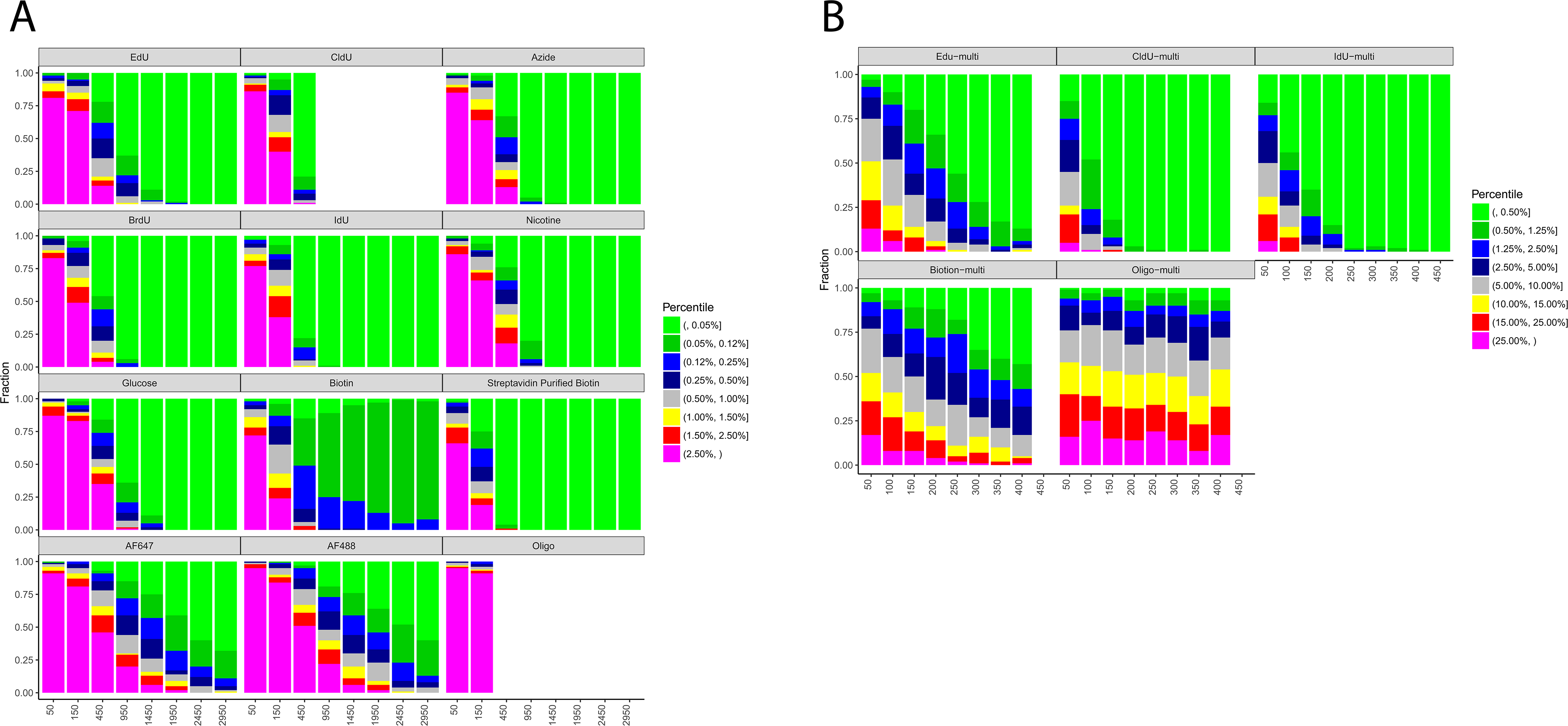
Sequencing depth required for the calling of base analogs in mixed samples with unmodified and modified reads. A). Modeling using single modified base templates. The X axis represents the number of reads used for the analysis, and the Y axis shows the fraction a modified site can be ranked by percentile among all sites for each sample. Analysis was performed using Stouffer’s combined statistics. B). Modeling using templates with multiple modifications for the modified region between 3042 and 3104 with tracks of thymidine analogs. IdU and CldU modifications cause the strongest signal shifts and require the least number of reads for reliable detection.

To determine the number of reads required to identify a region with multiple modifications, we performed the same simulation with 14 thymidine analogs in a single template (Fig. 5B). Multiple EdU modifications required 250 reads for reliable detection (percentile 0.05 or between the 0.05 and 0.12 percentiles). For other analogs CldU and IdU, only 100 reads proved enough for correct calling (percentile 0.05 or between the 0.05 and 0.12 percentiles). The template which contained multiple clicked biotins required 350 reads for accurate detection. This analysis showed that the presence of multiple modified bases in a DNA segment facilitates detection with NanoMod, requiring a smaller number of reads compared to a single modification. IdU was the modification that was most readily detectable in both the modelling of single modifications and multiple modifications.

### Detection of thymidine analogs incorporated in replicating mammalian DNA

To test if the MinION platform in conjunction with NanoMod can detect nucleotide analogs incorporated in replicating DNA, we incubated mouse embryonic stem (ES) cells from a pure C57BL/6J strain with 25uM IdU for 24h. Genomic DNA from unlabelled and IdU-incubated samples was then harvested, sheared in 6-8kb fragments and prepared for 1D sequencing on the MinION system. Error correction and signal annotation were performed with NanoMod. Base calling was accomplished with Albacore v2.3.1 and long reads were aligned with the GRCm38 (mm10) mouse reference genome. We aligned all reads from the IdU sample to the mouse reference genome and used a set of IdU-covered reads and control reads in the subsequent analysis. These reads were from different, rather than the same genomic location. Reads were split in 9-mers with no thymidines (VVVVVVVVV), V=AGC, 1 thymidine in the center of the k-mer (VVVVTVVVV) or 2 and more thymidines at different positions on the plus strand of the k-mer. This classification included 62 non-T k-mers and 277 T-containing k-mers of different sequence composition with coverage of 500 to 1500 reads per k-mer. Current signals were collected from the central T as well as 2 nucleotides upstream and downstream of the base and used in a Kolmogorov-Smirnov test to calculate p-values. Stouffer’s method was then applied to calculate a combined p-value for all 9-mers, and the result was plotted on a logarithmic scale. Comparing IdU labelled samples with unlabelled controls revealed that about 75% of T-containing k-mers in the IdU sample had combined log10 p-values between −308 and −3, while the percentiles of non-T k-mers were evenly distributed in their log10 P value, showing that IdU substitution in T-containing k-mers generates a detectable signal shift (Fig. 6A, B). Comparison of k-mers containing either 1T or 2T resulted in greater significance of detected differences: 80% of 1T and 2T-contaning 9-mers had a combined log10 p-values between −308 and −3 (Fig. 6C, D).

**Figure 6.**
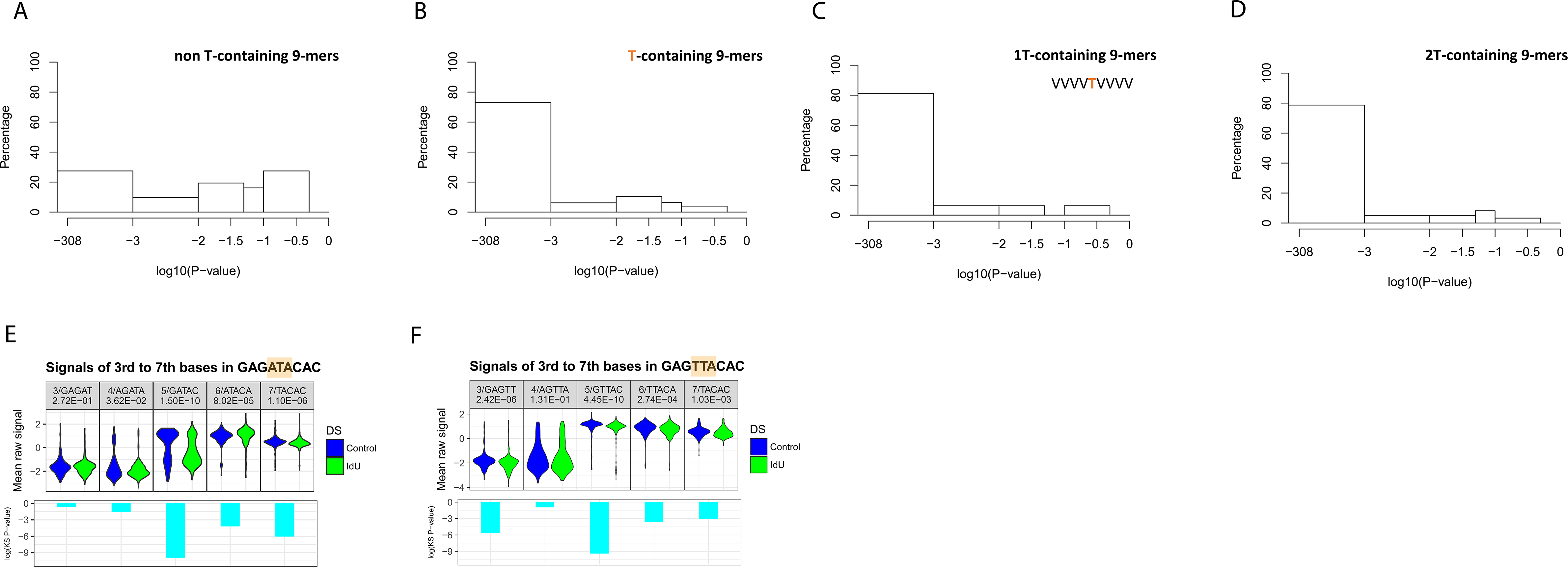
Detection of IdU incorporated into replicating mammalian DNA. A). Combined p-values calculated with Stouffer’s method for non-T containing 9-mers. The combined p-value includes p-values for the central base of the 9-mer as well as 2 bases upstream and downstream. B). Combined p-values for T-containing k-mers. All T-containing 9-mers were considered, including ones with multiple Ts. C). Combined p-values for 1T-contaning k-mers. This analysis included 9-mers with the thymidine base in the middle. D). Combined p-values for 2T-containg k-mers. The first T was set to the middle of the 9-mer and the other T could take any other possible position. E). Signals changes as a selected 9-mer from IdU-substituted genomic DNA with the sequence GAGATACAC, and F). GAGTTACAC was separated in 5 different k-mers. The k-mers were covered by 200-400 reads.

To determine whether signals from a k-mer of a specific sequence could be used to distinguish IdU from thymidine, two selected 9-mers GAGATACAC and GAGTTACAC were divided into 5 different k-mers. Electrical signals from control and IdU labelled samples of the same k-mer were plotted and compared using NanoMod. A significant shift was seen in the IdU containing k-mers, with the shift depending on the position of the T within the k-mer (Fig. 6E, F). This analysis used 427IdU/229control reads for GAGATACAC and 310IdU/211control for GAGTTACAC to identify an IdU containing k-mer in cellular DNA. This analysis shows that thymidine analogs, incorporated by replicating mammalian cells, can be called with nanopore sequencing, and that T containing k-mers of the same sequence, from different locations of the genome can be combined for the calling of base modifications. The combination of approximately 200 different k-mers on a single read should provide a tool to identify replicated DNA from long nanopore reads.

## DISCUSSION

Analysis of DNA replication at a single molecule level and genome would greatly increase our understanding of DNA replication and genetic stability in development and disease. Towards this aim, we show here that pore sequencing with the MinION can be used in conjunction with NanoMod to detect a panel of thymidine modifications in synthetic templates as well as IdU incorporated in DNA of replicating mammalian cells. Smaller analogs EdU, CldU, BrdU and IdU generated signals centered at the modified position and affecting 2-4 neighboring bases on each side with IdU causing the largest shift in pore current. In contrast, bulkier moieties, such as Biotin, generated widely spread signals affecting as many as 10 nucleotides in the neighborhood of the modified base.

Furthermore, we demonstrate that click cycloaddition reactions can be used to generate any desired chemical change after incorporation of a nucleotide with a reactive group. These analogs could be detected without further modification, and the clicking of bulky adducts again showed a wider spread in signals. Though click chemistry using copper catalysts can potentially introduce DNA damage, novel copper-free click chemistry is now available and will likely be more suitable for this application [19]. For example, AmdU is an azide group-containing analog, which is incorporated during DNA replication and reacts with alkynes in copper-free reactions, allowing the formation of a bulky adduct while protecting DNA from damage [20, 21]. A base with a vinyl-group, such as VdU, also allows the introduction of further modifications via copper-free alkene-tetrazine ligation[22].

The difference in signal spread between small analogs and larger bulky adducts can be potentially leveraged to distinguish between a small and a bulky analog which opens up the possibility of using the platform to determine replication patterns. For example, studies on the progression of DNA replication may use the sequential application of two analogs, such as IdU and EdU with the intent to modify the latter further with biotin and form a larger base. The use of two analogs is routinely applied in studies of fork stalling, origin mapping, analysis of replisome speed and direction. The ability to add information on location in the genome on single DNA strands through pore sequencing would greatly facilitate our understanding of the pattern of DNA synthesis in normal and abnormal cells.

To this end, we provide a proof of principle that noncanonical bases that can be incorporated into cells during DNA replication can be detected with nanopore sequencing. We chose IdU for analysis, as it performed best in modeling to detect a modified nucleotide when 20% of T’s are substituted. Using IdU labelling through a complete cell cycle, we demonstrate that IdU-substituted mammalian genomic DNA, can be distinguished from control DNA. A coverage of 200-300 per k-mer was sufficient to observe significant differences. As these k-mers are from different locations of the genome, k-mers from the same read may be combined for the analysis of labelled segments of a single DNA strand.

Applying this analysis to single reads will allow the assembly of detailed replication dynamics maps, which inform on origin usage and fork progression on single DNA strands. This is currently not possible with existing methods, which require amplification and library preparations, resulting in population averages. Therefore, sequencing DNA replication using nanopores, termed here Replipore sequencing, is a novel technology with significant potential in research and diagnostics.

Cell proliferation plays an important role in many diseases. Abnormalities in DNA replication can result in genome instability, cellular senescence or apoptosis, and are therefore highly relevant to regenerative medicine. Furthermore, DNA replication is cell type specific and dependent on epigenetic principles that also regulate cell type specific gene expression. Due to the fundamental requirement of DNA replication in cell cycle progression, the cell type specificity of the replication program may have an important role in determining limitations in cell proliferation [23]. Thus, insights about polymerase progression derived from sequencing modified nucleotides through nanopores will likely prove instrumental to our understanding, diagnosis and ultimately treatment of diseases characterized by cell cycle abnormalities.

## AVAILABILITY

NanoMod is an open source code available at https://github.com/WGLab/NanoMod.

## ACCESSION NUMBERS

The datasets generated and analyzed in this study will be available at the European Nucleotide Archive.

### ACKNOWLEDGEMENT

We thank Marcus Stoiber from the Lawrence Berkeley National Laboratory of Environmental Genomics and Systems Biology in Berkeley, CA for assistance in getting started on nanopore sequencing and base calling.

---

Supplementary Figure 1. Substitution of Mammalian Genomic DNA with EdU. A). Dot blots with EdU-substituted genomic DNA clicked with A647-azide. B). A647-fluorescence measurement of EdU-substituted DNA. The standard curve was prepared with products of PCR reactions ran with known concentrations of dTTP to dEdUTP and processed through click cycloaddition with A647 azide. The analysis shows ∼20% substitution in genomic DNA after 24h of incubation with 10uM EdU.

