## Supplementary figures and images for "Detection of Base Analogs Incorporated During DNA Replication by Nanopore Sequencing"

### Supplementary Figure 1

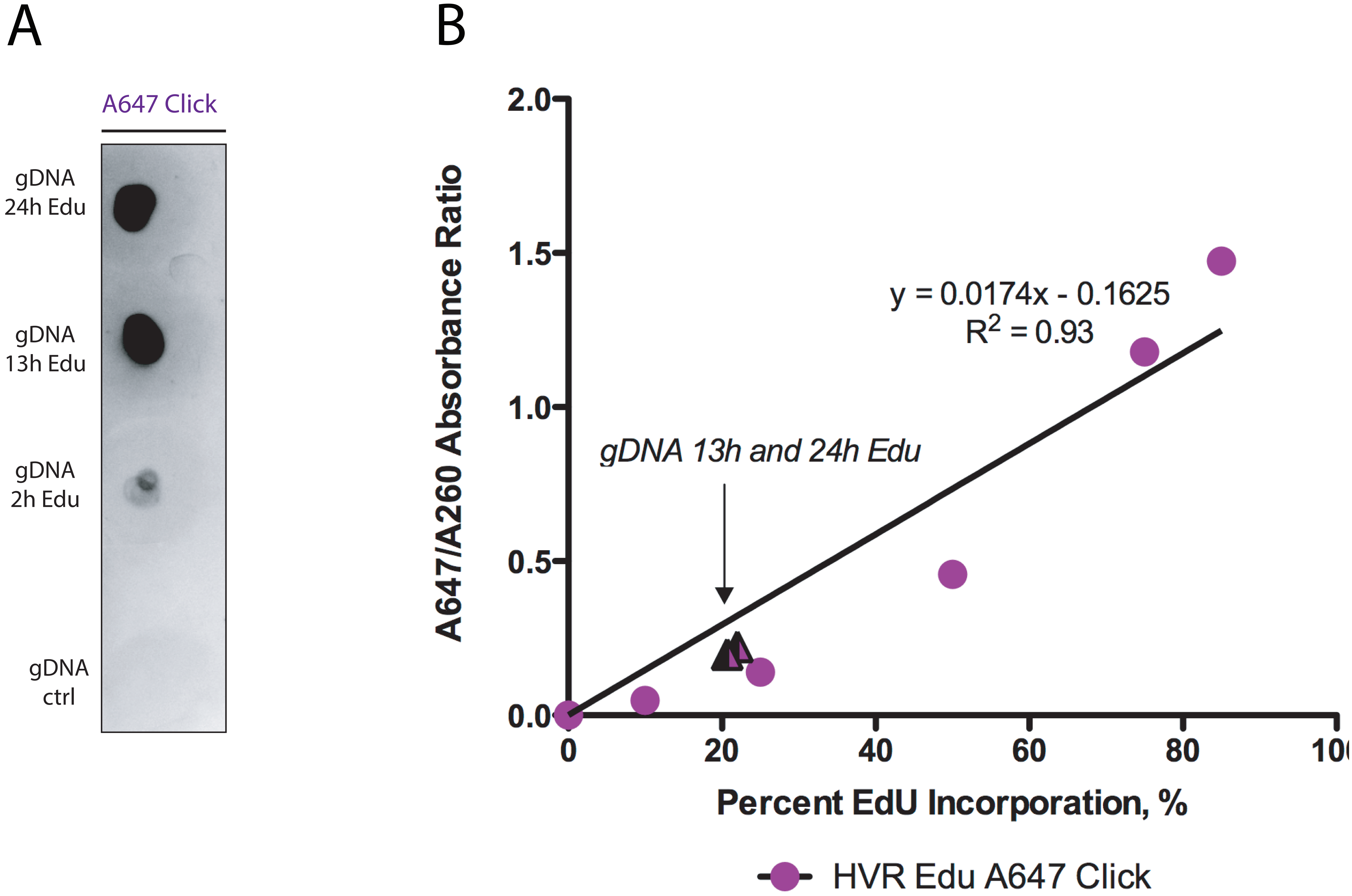
